# Processing of Reversed Replication Forks is Required for the Resolution of Replication-Transcription Conflicts

**DOI:** 10.64898/2026.03.20.713164

**Authors:** Juan Carvajal-Garcia, Houra Merrikh

**Affiliations:** Department of Biochemistry, Vanderbilt University School of Medicine, Nashville, TN 37232, USA

## Abstract

DNA replication and transcription occur simultaneously on the same template, leading to conflicts between the two machineries. Conflicts stall replication forks, lead to genome instability, mutagenesis, and the formation of deleterious R-loop structures.

There are two types of conflicts: depending on the strand where a gene is encoded, the two machineries either meet head-on or co-directionally. The adverse outcomes of conflicts in the head-on orientation are significantly more detrimental to cells compared to co-directional conflicts. Despite many studies across various organisms, how the replication fork structure is impacted by these encounters remains unclear. Here, we performed an unbiased genetic screen using a transposon library to identify factors essential for surviving head-on conflicts in *Bacillus subtilis*. Our screen identified three hits: RNase HIII, AddA and AddB. Our prior work had shown that RNase HIII, which processes R-loops, is essential for surviving head-on conflicts. However, AddA and AddB, which function together as a complex to process blunt DNA ends, have not been previously identified as *essential* conflict resolution factors. Through follow-up genetic and biochemical analyses, we found that the helicase activity but not the nuclease activity of this complex is required for conflict resolution. Based on the fundamental properties of DNA at replication fork structures, our work collectively indicates that upon head-on conflicts, the nascent strands form a reversed fork structure, which is then unwound by AddAB, which leads to re-annealing to the parental strands. This process re-establishes an intact replication fork that can be used to restart replication after conflicts with transcription.

## Introduction

DNA replication and transcription share a common template and can occur simultaneously, leading to conflicts between the replication and the transcription machineries. These conflicts occur either co-directionally when a gene is encoded on the leading strand, or head-on when a gene is encoded on the lagging strand. While both types of conflicts can be detrimental for cells and lead to genome instability (1), head-on conflicts have been shown to have more adverse effects, which include replication stalling (2–4), replication restart (3), chromosome breaks (5, 6), and higher mutation rates (7, 8). Accordingly, across highly divergent bacteria, most genes are encoded on the leading strand, and this is especially true for those that are highly expressed (9); meanwhile, in human cells, origins of replication tend to coincide with transcription start sites, which has been proposed as a way to ensure co-directionality of replication and transcription, at least for protein coding genes (10).

However, both bacteria and humans have plenty of genes in the head-on orientation, which require specialized cellular pathways that help resolve head-on replication-transcription conflicts. For instance, head-on conflicts lead to the preferential formation of R-loops (5, 11), three stranded nucleic acid structures that form when the nascent RNA reanneals to its template DNA and displaces the non-template DNA strand. R-loops can lead to genome instability, mutagenesis, replication stress, and chromosomal breaks, and need to be resolved for proper cellular function (5, 11, 12). Other mechanisms that promote genome stability at head-on conflicts are homologous recombination (4, 13), transcription-coupled repair (14, 15), resolution of topological stress (16), and coupling between transcription and mRNA processing (17).

Using the Gram-positive model bacterium *Bacillus subtilis*, we set out to determine what other factors prevent cell death due to head-on conflicts. To address this question, we used a transposon mutagenesis screen. Our screen revealed three genes that promote cell survival specifically in cells containing an engineered head-on conflict: *rnhC*, *addA*, and *addB*. While the role of *rnhC*, which codes for RNase HIII, in head-on conflict resolution is well understood (11), an essential role of *addA*, and *addB*, which generate the AddAB heterodimer complex, in conflict resolution has not been previously described.

AddAB is a helicase-nuclease that is involved in the first step of homologous recombination in Gram-positive bacteria, and is a functional homolog of the *Escherichia coli* RecBCD helicase-nuclease complex (18). AddAB recognizes a blunt or nearly blunt double stranded DNA end (19). Our data show that, in the context of head-on replication-transcription conflicts, these DNA ends are generated by replication fork reversal. Reversed replication forks are four-way junctions that are formed across the tree of life when replication is perturbed, such as by uncoupling of the leading and the lagging strands (20). These reversed forks need to be restored, either by unwinding or by degradation. Here, we determined that the helicase but not the nuclease activity of the AddAB complex is required for processing of the stalled replication forks post-conflicts. Taken together, our results show that fork remodeling, specifically re-annealing of the nascent strands, is a key process for continuing DNA replication when it stalls due to transcription.

## Results

### A screen to find genes required for replication-transcription conflict resolution

We set out to find genes that are involved in the resolution of replication-transcription conflicts by employing a transposon mutagenesis screen. We generated transposon libraries in cells where we inserted the *luxABCDE* operon in the head-on orientation to replication, into the *amyE* locus, under the control of the inducible promoter *Pspank(hy)*. We have shown in the past that when this promoter is turned on by the addition of isopropyl β-D-1-thiogalactopyranoside (IPTG), conflicts between transcription of the *luxABCDE* operon and replication lead to cell death in the absence of conflict resolution factors (11). This method cannot identify essential genes, such as the helicase PcrA, which mitigates replication transcription conflicts (21).

To generate the transposon library, we used a plasmid harboring a hyperactive mariner-Himar1 transposase and a kanamycin resistance gene flaked by the invert repeats that this transposase recognizes. Because this plasmid has a temperature-sensitive origin of replication in *B. subtilis*, once the transposon has been mobilized to the chromosome in the permissive temperature, the rest of the plasmids can be eliminated by growing the cells in the restrictive temperature (see Methods for more details) (22, 23). The transposon library was then grown in parallel, in cultures without IPTG (conflict OFF) or with IPTG (conflict ON), followed by DNA extraction and sequencing of the chromosomal regions adjacent to the transposon insertion (Fig 1A).

**Figure 1.**
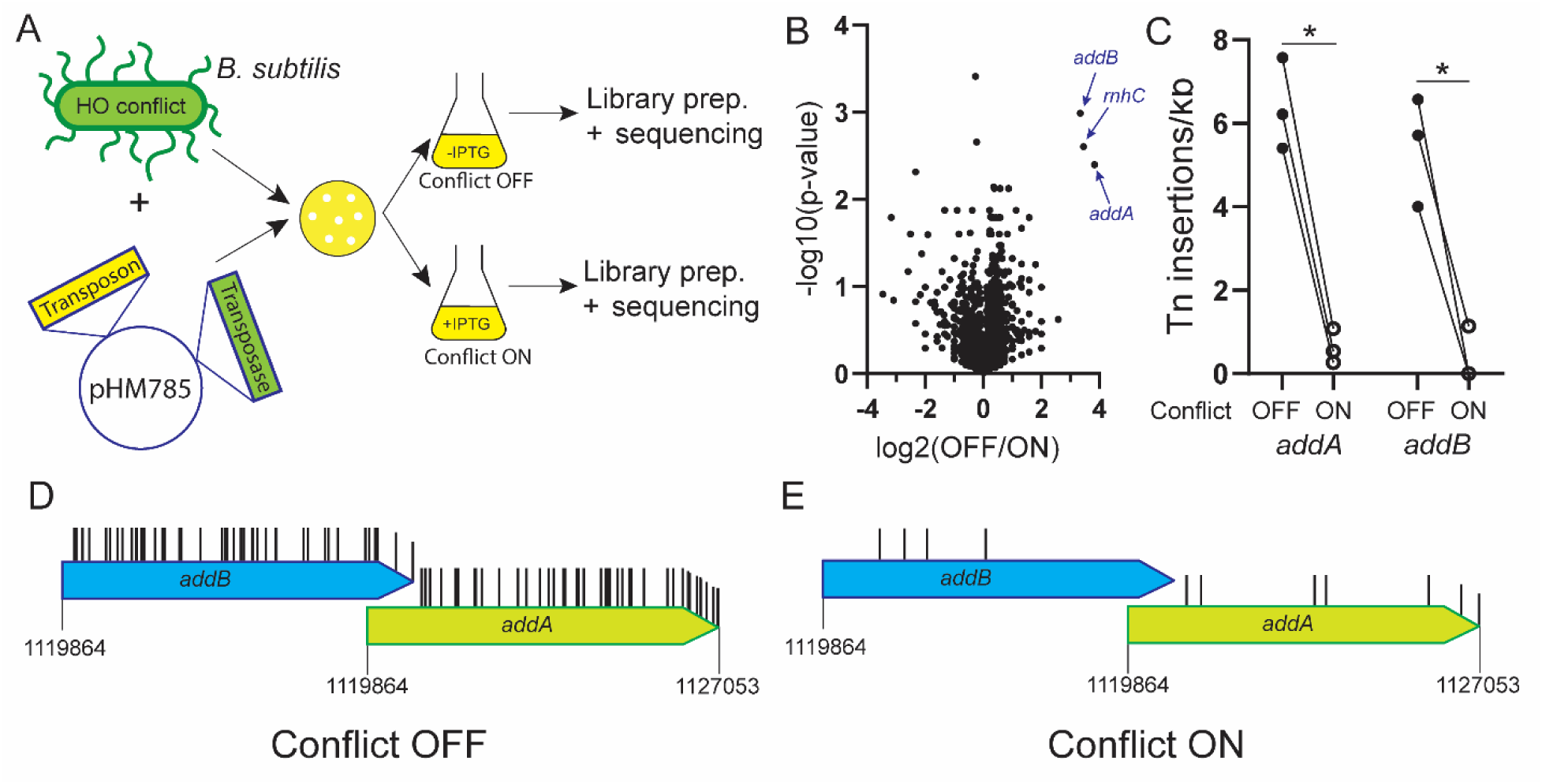
A transposon mutagenesis screen identifies AddAB as a head-on replication-transcription conflict resolution factor. A) Schematic of the transposon screen. A whole-genome transposon library was made in cells expressing an inducible gene in the head-on orientation, and this library was grown in parallel in media without (Conflict OFF) or with 1 mM IPTG (Conflict ON), followed by high throughput sequencing of the transposons. B) Volcano plot showing the log_2_ of the ratio of transposon insertions per gene (Conflict OFF/ON) on the X axis and the -log_10_ of the *p*-value (two-tailed t-test) on the Y axis. The three main hits (*addA*, *addB*, and *rnhC*) are highlighted. C) Number of transposon insertions per kilobase of DNA found in *addA* and *addB* in the conflict OFF and ON condition. D) Location of the transposon insertions found in *addA* and *addB* in the conflict OFF and ON condition. Statistical significance was assessed by 2-way ANOVA with multiple comparisons. \**P* < 0.05.

We determined the number of transposon insertions per coding sequence by aligning the sequence reads to the *B. subtilis* genome and obtained an average of approximately 11 transposon insertions per transcription unit. We plotted the resulting data in a volcano plot, with the log_2_ of the ratio of transposon insertions per gene when the conflict is OFF over the transposon insertions per gene when the conflict is ON on the X axis, and the -log_10_ of the p-value (two-tailed, unpaired t-test) on the Y axis (Fig. 1B). This plot revealed three genes that were strongly depleted specifically when the conflict in on: *rnhC*, *addA* and *addB* (Fig. 1B-D).

We previously described that the role of *rnhC*, which encodes for the RNase HIII enzyme is essential in head-on conflict resolution (11), confirming that our screen was highly efficient and can identify essential conflict resolution factors efficiently. However, since *addA* and *addB* have not been described as essential conflict resolution factors, we decided to follow up these hits. These two genes encode for the proteins AddA and AddB, which form a heterodimer and can only function when they are in a complex together (24).

### The AddAB complex is required for survival during head-on conflicts

To validate the results of our screen, we plated serial dilutions of WT cells and cells with either the *addA* or *addB* gene deleted, on plates that either do not induce the conflict (without IPTG, conflict OFF) or do induce the conflict (with 1 mM IPTG, conflict ON). While without induction, we observed a small decrease in the viability of the cells (Fig. 2A), inducing the conflict led to a large decrease in survival (>90%) specifically in Δ*addA* and Δ*addB* cells (Fig. 2A-C).

**Figure 2.**
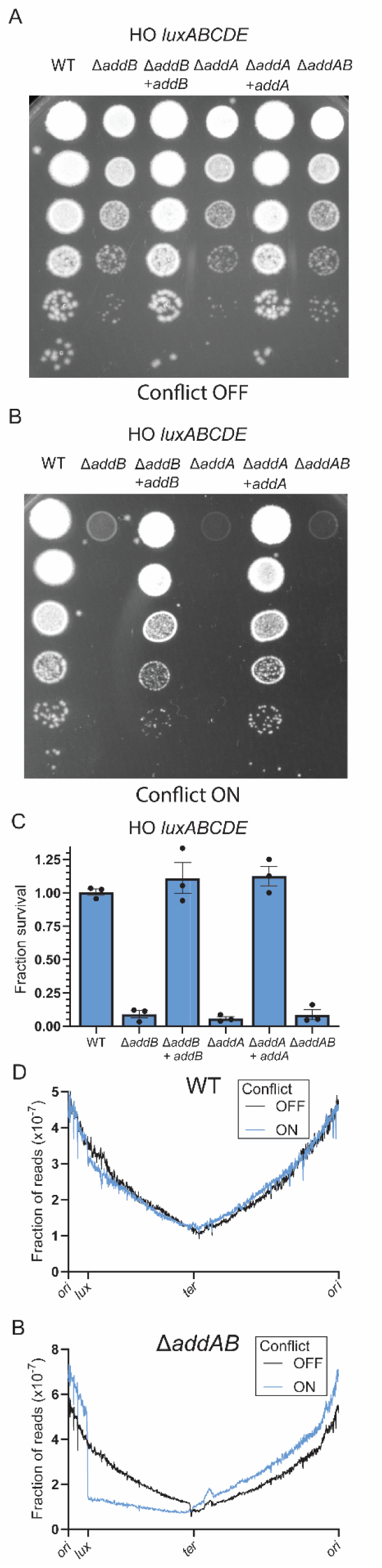
AddAB are required for cell viability in cells experiencing a severe head-on conflict. A-B) Serial dilutions of cells of the indicated genotype with the *luxABCDE* conflict integrated into *amyE* in the head-on orientation plated in a plate without IPTG (A) or with 1 mM IPTG (B). C). Fraction of surviving cells of the indicated genotype with the *luxABCDE* conflict integrated into *amyE* in the head-on orientation calculated as the ratio of the number of colonies when the conflict is ON over the number of colonies when the conflict is OFF. D-E) Fraction of reads that map to the indicated coordinates (5 kb window) of the *B. subtilis* genome, smoothed using a second other polynomial using 10 neighbors on each side in WT (D) or Δ*addAB* (E) cells growing exponentially when the *luxABCDE* conflict is OFF (no IPTG) or ON (1 mM IPTG). The

Importantly, expression of the WT version of each gene, respectively, from a different chromosomal location (*thrC* locus) fully rescued the viability of the cells when the conflict was on (Fig. 2A-C). Moreover, cell lacking both *addA* and *addB* (Δ*addAB*) had a similar loss in viability as the single mutants (Fig. 2A-C). This indicates that, as expected and based on the literature, AddA and AddB work together to promote cell viability in the presence of severe head-on replication-transcription conflicts.

To control for sequence bias and genomic location, we repeated these experiments with a different gene, *lacZ*. We integrated the *lacZ* gene in the head-on orientation and under the control of the same promoter as the *luxABCDE* conflict, into the *thrC* locus. We observed a similar phenotype, albeit milder (Fig. S1A-B). This is consistent with previous data indicating that conflicts within the *luxABCDE* operon are more severe than the *lacZ* gene (11). The *lux* operon is roughly 5.5 kb long whereas *lacZ* is 3 kb long. By definition, the longer the gene, the more conflicts can occur, which explains the difference in the severity between these two engineered conflicts. As an additional control, we tested the role of AddAB in the resolution of co-directional conflicts using *luxABCDE* and observed no difference in viability between WT and Δ*addAB* (Fig. S1C-D). This result is expected given that co-directional conflicts are readily resolved and are generally not nearly as severe as head-on conflicts (3). The absence of a phenotype different from WT with the co-directional conflict is either because we do not have the resolution to detect a role for AddAB at these regions, or that AddAB specifically resolves head-on conflicts.

Head-on conflicts have been shown to completely stall replication in cells lacking conflict resolution factors (6, 16). Therefore, we determined the genome copy number to test whether this is also the case for Δ*addAB* mutans. To do this, we used high throughput sequencing of the genomes from either exponentially growing WT or Δ*addAB* cells, containing the engineered *luxABCDE* operon, under conditions where the gene is either not induced (conflict OFF) or is induced (conflict ON), after which we aligned them to the *B. subtilis* genome. Because *B. subtilis* has one defined origin of replication per chromosome, the sequences surrounding the origin have more reads that align to them, with the number of reads progressively decreasing along each chromosome arm, to a minimum that overlaps with the terminus. Moreover, due to muti-fork replication, a phenomenon observed when bacteria grow in rich media, the ratio of reads that align to the origin and reads that align to the terminus is higher than 2 (approximately 4.5 in our case) (Fig 2D-E). In WT cells, when the conflict was on, we observed a small decrease in sequencing reads right after the conflict region, indicating a mild slowdown of the replication fork (Fig. 2D). However, in Δ*addAB* cells, upon induction of the conflict, we observed a sharp decrease in the number of reads aligning to the conflict region, which continues to the terminus (Fig. 2E). This indicates that in the absence of AddAB, replication is unable to continue past the conflict region, which by definition will lead to the cell death as described above.

### The role of AddAB in conflict resolution is independent of R-loop formation

In addition to *addA* and *addB*, we had another hit in the screen, *rnhC*, which codes for RNase HIII. We have established in the past that R-loop processing by RNase HIII underlies its requirement for survival upon severe head-on replication-transcription conflicts (5, 11). We set out to determine if AddAB and RNase HIII function in the same genetic pathway during conflict resolution. If so, this would suggest that AddAB might be involved in R-loop formation or processing. This experiment required the generation of an *addA addB rnhC* triple mutant.

However, we were unable to make this strain, suggesting that *addAB* might be synthetic lethal with *rnhC*, as is the case with other genes involved in homologous recombination, such as *recA* (25). We confirmed this synthetic lethal interaction by using a conditional degron system in which expression of the SspB protein targets any protein with a C-terminus SsrA tag for ClpXP-dependent degradation (26). We added the SsrA tag to the C-terminus of RNase HIII, in a strain that expresses SspB under the control of the *Pspank* promoter, which leads to RNase HIII degradation upon induction of *sspB* expression (Fig. S1A). Consistent with *rnhC* and *addAB* being synthetic lethal, we observed no surviving colonies when we turned on the expression of *sspB* with IPTG (Fig. S1B-C).

To determine whether AddAB and RNase HIII are in the same genetic pathway during conflict resolution, we incorporated the *luxABCDE* conflict into this system, and induced the expression of both the *luxABCDE* operon and the *sspB* gene with 100 µM IPTG. Even with reduced *luxABCDE* expression, we still observed a strong decrease in survival of the Δ*addAB* cells that have WT levels of RNase HIII (Fig. S1D-E; second column), yet smaller than when using 1 mM IPTG (70% lethality vs. >90% in Fig. 2C). When RNase HIII levels are decreased by induction of *sspB* in otherwise WT cells, we also observed a large yet not complete decrease in viability (Fig. S1D-E; third column). These intermediate phenotypes allowed us to observe an additive effect of decreasing RNase HIII levels in Δ*addAB* cells, as we did not detect any survival in Δ*addAB* cells with decreased levels of RNase HIII and the head-on conflict (Fig. S1D-E; fourth column). To confirm that these cells are dying due to the head-on conflict and not due to the synthetic lethality we observed between these three genes, we used the *luxABCDE* operon in the co-directional orientation. We observed that Δ*addAB* cells with decreased levels of RNase HIII and a co-directional conflict are able to survive in 100 µM IPTG (Fig. S1D-E; fifth column), indicating that the reduced levels of RNase HIII protein are enough to sustain viability in Δ*addAB* cells in the absence of a head-on conflict. This indicates that AddAB and RNase HIII are in different genetic pathways when cells must resolve a severe head-on conflict. Additionally, overexpression of RNase HIII, which removes R-loops at head-on conflict regions (11), does not rescue the viability of Δ*addAB* cells upon induction of the conflict (Fig. S1F-G). This result also supports that AddAB does not have a role in R-loop formation or processing.

### AddAB’s role in conflict resolution is independent of double strand break repair

AddAB’s best understood function is in double strand break repair. In the beginning of homologous recombination, it generates the 3’ ssDNA tails that allow RecA, the main protein involved in homologous recombination in bacteria, to load into the DNA and perform its strand exchange function (18). Because of this, and even though RecA did not come out as a significant hit in the screen (Fig. 3A), we tested the role of RecA in conflict resolution. In agreement with our screen, we observed no difference in survival between WT and Δ*recA* cells when the *luxABCDE* conflict was on (Fig. 3B). Moreover, the replication profile of Δ*recA* cells with the conflict on showed a similar mild replication slowdown as in WT cells, not the complete replication stalling that we observed in Δ*addAB* cells (Fig. 3C). This suggests that the role of AddAB in conflict resolution is not related to homologous recombination.

**Figure 3.**
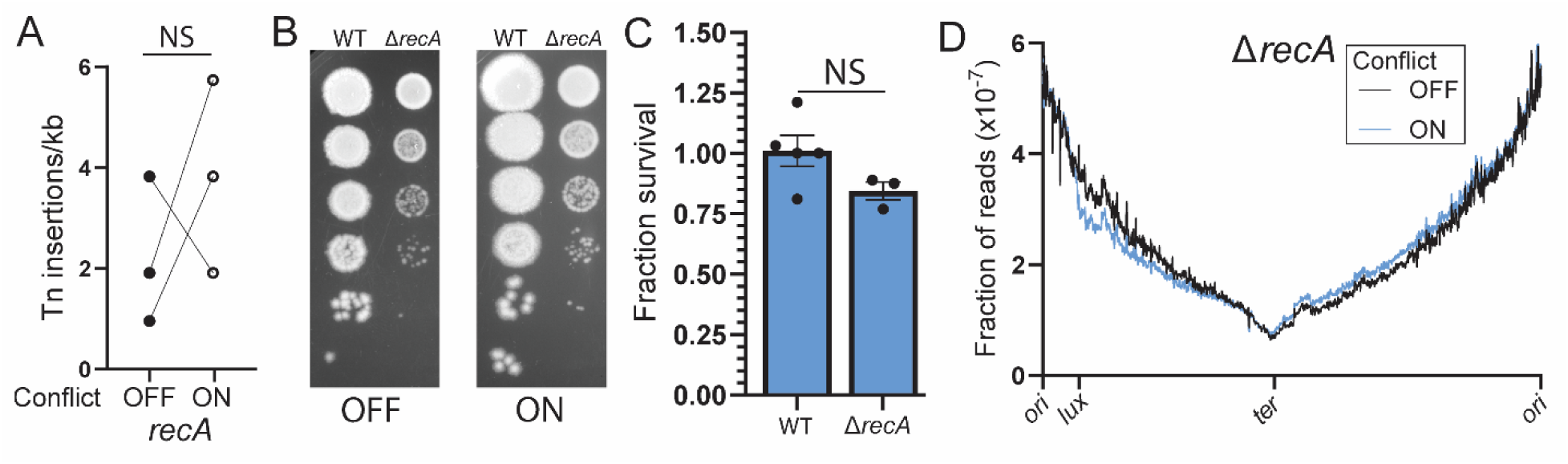
RecA is not involved in the resolution of head-on replication-transcription conflicts. A) Number of transposon insertions per kilobase of DNA found in *recA* in the conflict OFF and ON conditions. B-C) Survival assay of cells of the indicated genotype with the *luxABCDE* conflict integrated into *amyE* in the head-on orientation. Fraction survival was calculated as in Fig. 2C. D) Fraction of reads that map to the indicated coordinates of the *B. subtilis* genome in Δ*recA* cells growing exponentially when the *luxABCDE* conflict is OFF (no IPTG) or ON (1 mM IPTG). Statistical significance was assessed by an unpaired t-test. NS: *P* < 0.05.

Because *B. subtilis* can repair double strand breaks by non-homologous end joining, we reasoned that perhaps this pathway was masking RecA’s role in conflict resolution. To test this, we introduced the *luxABCDE* conflict into cells lacking the main proteins known to be involved in this pathway in *B. subtilis*: Ku, which is involved in DNA-end-binding, and LigD, the double strand DNA ligase (27), and made double mutants with RecA. We did not observe a decrease in cell viability when the conflict was on in neither single nor double mutants (Fig. S2A-B). This is consistent with previous findings showing that, in *B. subtilis*, the non-homologous end joining pathway is restricted to spores (27). This indicates that these pathways are not required for conflict resolution.

Additionally, the functional homolog of AddAB in *E. coli*, RecBCD, has been shown to be involved in replication termination, through a mechanism that is independent of RecA and dependent of the nuclease complex SbcCD (28). To rule out that an analogous mechanism is responsible for AddAB-dependent conflict resolution, we tested the role of SbcC and SbcD in conflict resolution and did not observe any decrease in viability (Fig. S3C-D), consistent with the fact that these genes also did not come up in the screen.

### AddAB can degrade reversed replication forks

To initiate DNA degradation, AddAB requires a blunt or almost-blunt DNA end, as it is not able to degrade ssDNA that is bound by single-stranded binding protein (SSB) in vitro (19). The lack of a role for RecA in conflict resolution led us to hypothesize that AddAB might be acting in reversed replication forks, four-way junctions that are formed by annealing the nascent strands and re-annealing the parental strands. These structures have been observed in many organisms under conditions that lead to replication stress and provide a double stranded DNA end that AddAB can act on (20). Moreover, reversed forks have been observed by electron microscopy in *B. subtilis* cells in which the inducible *lacZ* conflict had been integrated into the *amyE* locus in the head-on orientation (29). Additionally, the genetic requirement for the resection machinery but not RecA in head-on conflict resolution has been observed in *E. coli*, and the formation of reversed replication forks has been proposed to explain this (6).

In this situation, AddAB could be degrading the annealed nascent strands, which allows for the fork to be restored. However, it is unknown if AddAB can perform this reaction. To test this, we expressed and purified *B. subtilis* AddAB (Fig. S4A). First, we confirmed that, as has been previously shown, purified AddAB can digest dsDNA but not ssDNA when SSB is included in the reaction (Fig. S4B-C) (19, 30), and that it requires ATP for digestion (Fig. S4D) (30). Then, we assessed its ability to digest the two reversed fork structures that can arise from a lagging strand replication block, as is the case in a head-on conflict. One of these has a short 3’ overhang after nascent strand annealing (Model 1), while the other one is blunt (Model 2) (Fig. 4A). To monitor the degradation of both nascent strands, we labeled one with the Cy5 and the other one with Cy3. When we incubate these substrates with AddAB, in the presence of ATP and SSB, we observed that it is able to degrade both annealed nascent strands (Fig. 4B-C, top bands). Accordingly, we observed the corresponding appearance of a band that represents digestion up until the junction (Fig. 4A-B, 29 nt band), as well as a smaller band (20 nt) that suggests some digestion past the junction. These results show that, at least in vitro, AddAB can restore reversed replication forks by nascent strand degradation.

**Figure 4.**
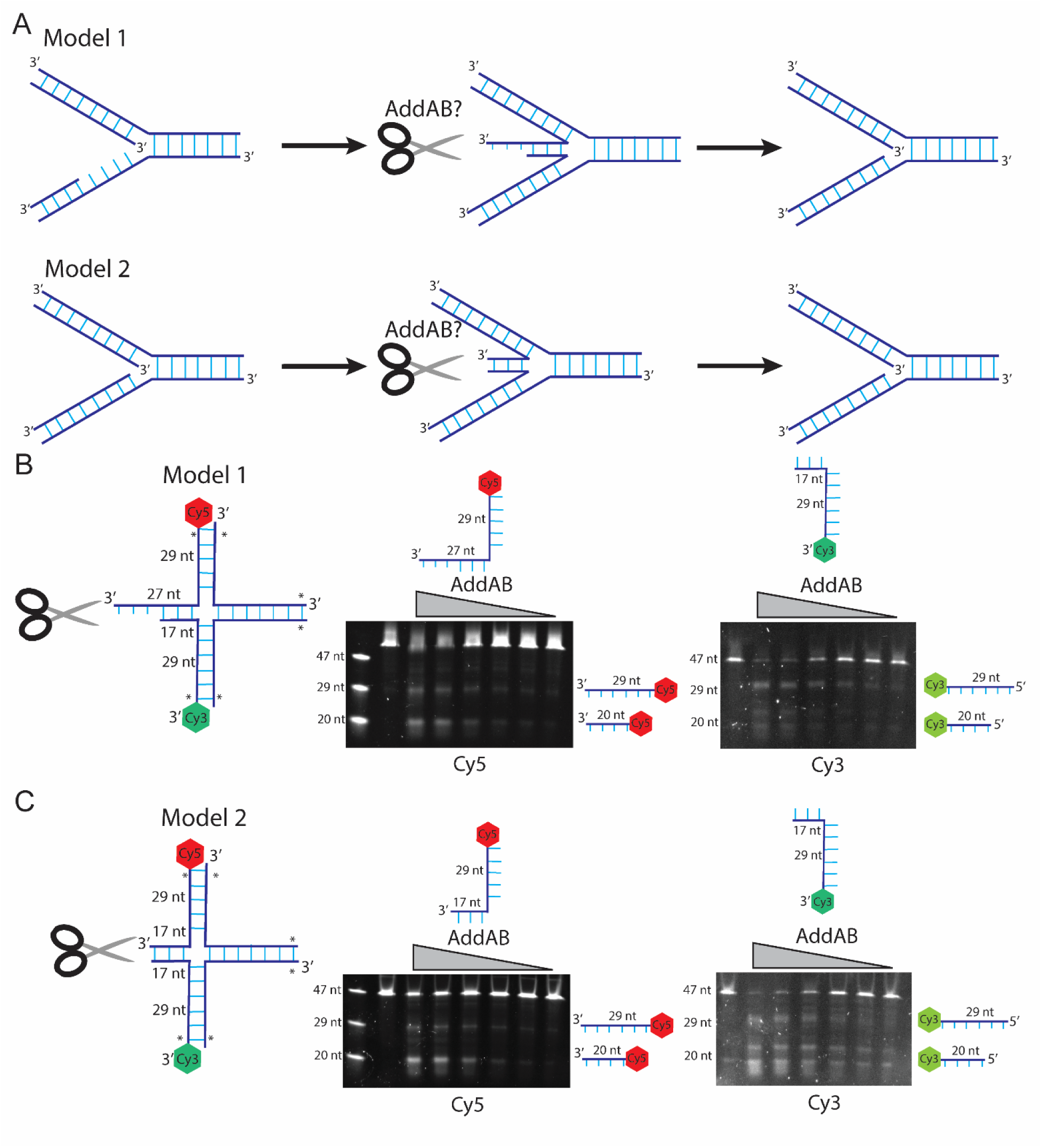
AddAB can degrade reversed replication forks. A) Schematic of the two types of four-way junctions that can be formed by a lagging strand block. B-C) 10% Urea gels of the indicated substrates after digestion with AddAB. The highest concentration of protein is 150 nM and the lanes to the right are 2-fold dilutions. Only the branch closest to the scissors is digestible, as the others have 3 phosphorothioate bonds in the DNA ends (asterisks). The labeled DNA oligos after DNA denaturation are shown above the gel. The pictures in the same panels are from the same gel, using either the Cy5 filter (left) or the Cy3 filter (right).

### The AddAB helicase but not the nuclease activity is essential for conflict resolution

To determine the precise mechanism behind AddAB conflict resolution in vivo, we set out to test which AddAB biochemical activities are involved in this process. In vitro, AddAB has three activities: AddA provides ATP dependent 3’ to 5’ helicase activity and 3’ exonuclease activity, and AddB provides 5’ exonuclease activity (Fig. 4A) (30). Combined, these activities lead to double stranded DNA degradation that is initiated by DNA unwinding, as a helicase deficient AddA is unable to degrade DNA (Fig. 4A) (30). Upon encountering the *B. subtilis* Chi site (5’-AGCGG-3’) (31), the 3’ exonuclease activity of AddA is suppressed, rendering a 3’ ssDNA tail as the product of the reaction (Fig. 4A) (30).

We complemented Δ*addA* or Δ*addB B. subtilis* strains with *addA* and *addB* separation of function mutants that specifically lack one biochemical activity: AddB^N^ (D961A) which lacks 5’ exonuclease activity, AddA^N^ (D1172A) which lacks 3’ exonuclease activity, and AddA^H^ (K36A) which lacks helicase activity (Fig. 5A) (30). We then introduced the mutants into cells carrying the *luxABCDE* head-on conflict. We observed that the 5’ exonuclease activity of AddA is dispensable for conflict resolution, that the 3’ exonuclease activity of AddB is partially required, and that the helicase activity of AddA is absolutely requited (Fig. 4 B-C).

**Figure 5.**
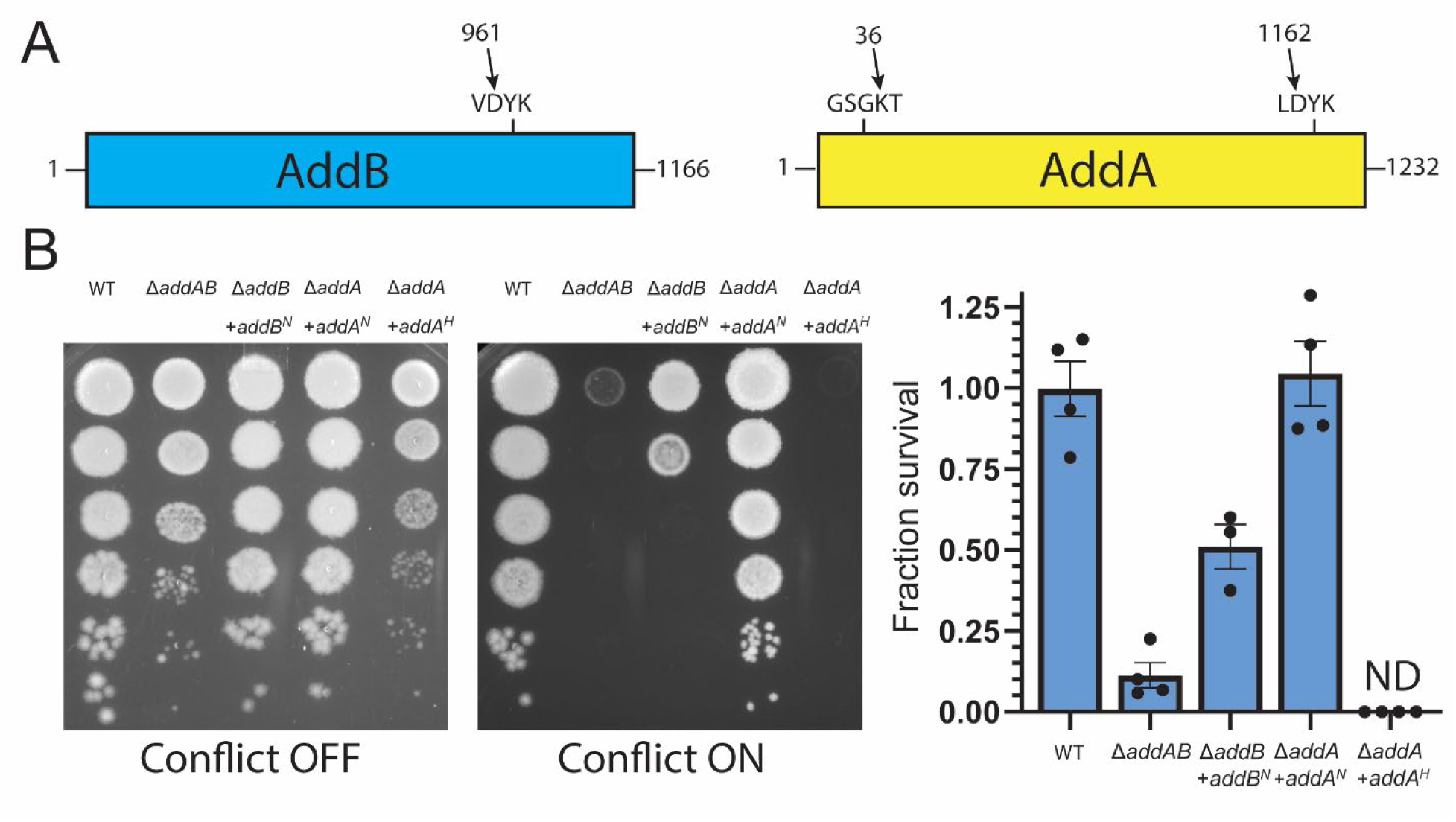
The AddAB helicase activity is essential for head-on conflict resolution. A) Schematic of AddB and AddA with the nucleases and Walker A (AddA) motifs highlighted. The mutated amino acids to create the separation of function mutants are indicted. B) Survival assay of cells of the indicated genotype with the *luxABCDE* conflict integrated into *amyE* in the head-on orientation. Fraction survival was calculated as in Fig. 2C.

These data suggest that while AddAB can digest reversed forks, its main mechanism of fork restoration might be to facilitate re-annealing of the nascent strands (Fig. 6, left). Such a mechanism would be analogous to Human RECQ1-dependent fork restoration (32). Meanwhile, nascent strand degradation might be a minor one (Fig. 6, right). Alternatively, the helicase activity of AddAB can be required for unwinding the annealed nascent strands, which would allow other ssDNA exonucleases in the cell to digest them (Fig. 6).

**Figure 6.**
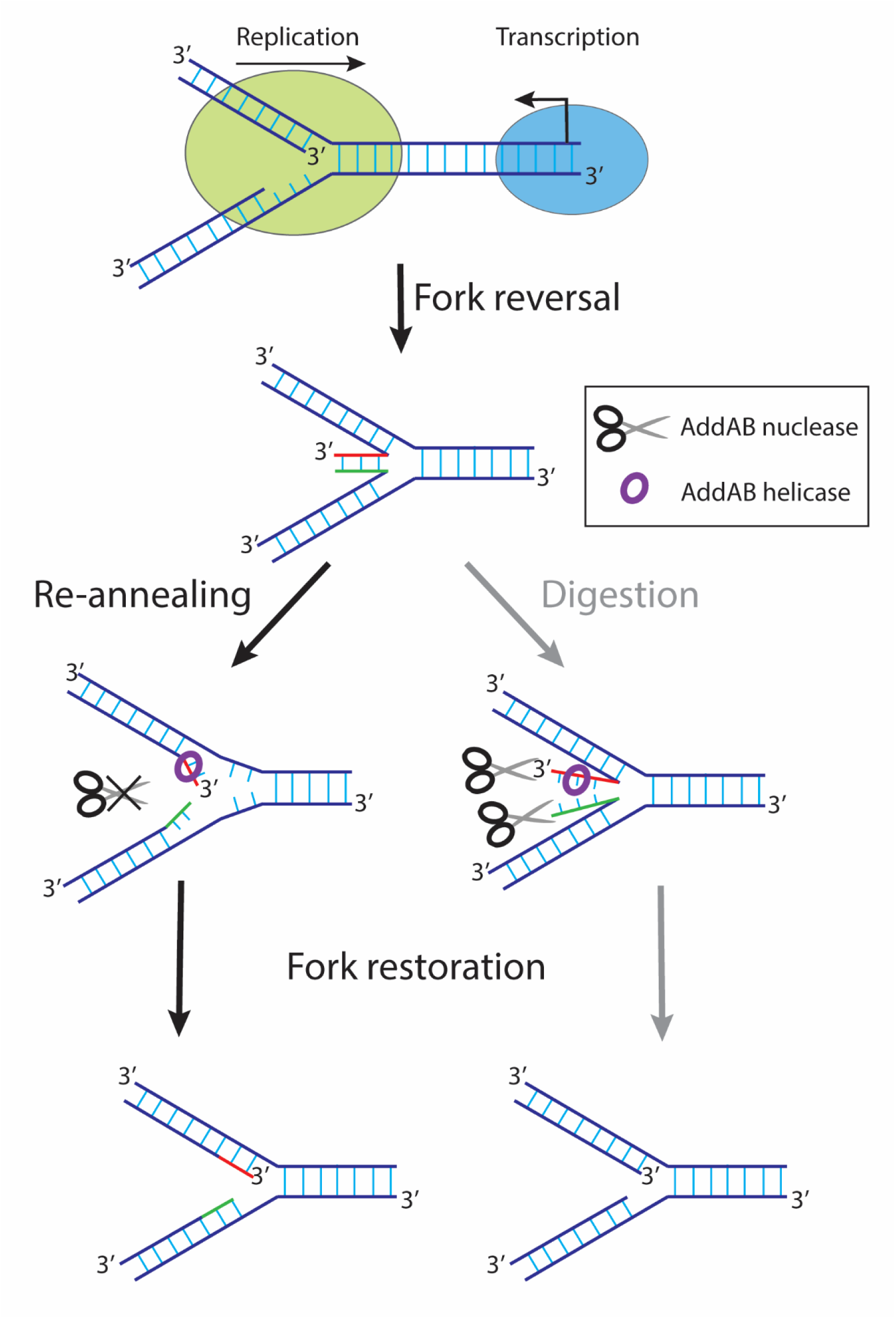
Schematic of the proposed model for replication-transcription conflict resolution by replication fork reversal. Upon encountering a highly expressed gene in the head-on orientation, the replication fork reverses, and AddAB can be restored in two ways. On the left, the helicase activity may re-anneal the nascent strands. On the right, the helicase and nucleases activities co-operate to degrade the nascent strands. Both mechanisms restore replication forks and allow replication to continue.

## Discussion

Transcription is a major threat to genome instability (33), and one of the main reasons for this is the fact that replication and transcription, two fundamental cellular processes, co-occur in time and space, leading to conflicts between the two machineries. These conflicts are detrimental for cells in both prokaryotes and eukaryotes. Therefore, cells have pathways that resolve these conflicts to promote genome stability and survival (1).

In this study, we set out to identify genes that are essential for the resolution of replication-transcription conflicts using an unbiased approach, a whole genome transposon mutagenesis screen. The results of this screen identified *rnhC*, *addA*, and *addB* as genes that encode for proteins necessary for conflict resolution (Fig. 1). We previously described the role of *rnhC*, which encodes for RNase HIII, in promoting survival of cells experiencing a severe head-on conflict. This finding in essence served as a positive control for the screening system we constructed. AddAB on the other hand had not been previously shown to be essential for conflict resolution. Our follow-up studies confirmed that AddAB is indeed required for survival in cells expressing the *luxABCDE* operon in the head-on orientation, which creates severe conflicts (Fig. 2A-C). Our genome copy number experiments showed that this is due to replication not being able to progress past the conflict region (Fig. 2D-E).

AddAB is a helicase-nuclease complex best known for carrying out the first step of homologous recombination. Surprisingly, *recA* was not identified in our screen as an essential conflict resolution factor (Fig. 3A), and we confirmed this in follow-up experiments (Fig. 3B-D).

Therefore, the role of AddAB in conflict resolution is unlikely to depend on its function in homologous recombination. Similarly, the other double strand break repair pathway known in *B. subtilis*, non-homologous end joining, was also not required for conflict resolution.

In vitro, AddAB requires a blunt or almost blunt DNA end to function, and we reasoned that the only process that can generate this type of DNA end, and be repaired independently of homologous recombination or non-homologous end joining, is replication fork reversal (6).

Importantly, reversed replication forks have been observed following replication stalling by transcription in *B. subtilis* cells (29). Approximately 10% of replication forks are reversed in WT *B. subtilis* cells experiencing a head-on conflict. This number increases to over 20% in Δ*rnhC* cells, supporting our conclusion that AddAB and RNase HIII are in different genetic pathways (Fig. S2). It is important to point out that the cells used for the experiments where reversed forks were observed contained AddAB, therefore the low number of cells with these structures is expected.

While the activities of AddAB are well established in vitro (19, 30), it remained unknown if this complex could degrade a reversed fork, leading to restoration of replication forks so they can continue replication. To test this, we purified *B. subtilis* AddAB, created the DNA structures most likely to arise from fork reversal at a lagging strand block, and showed that AddAB can degrade the annealed nascent strands, up until the junction or shortly past it (Fig. 4).

Strikingly, when we tested the role of the different biochemical activities of AddAB in promoting cell survival upon induction of the conflict, we observed that the helicase activity of AddAB is absolutely required for survival, yet the nuclease activities are not. The 5’ exonuclease has a partial role while the 3’ exonuclease is dispensable. This result led us to propose a model in which AddAB’s role in reversed fork restoration is through re-annealing of the nascent strands (Fig. 6, left pathway) and that degradation of the nascent strands might be less favored (Fig. 6, right pathway). Alternatively, the helicase activity might be promoting the unwinding of the nascent strands, that can then be degraded by other exonucleases in the cell. Interestingly, a recent study observed a similar requirement for only the helicase activity of AddAB and not the exonucleases in the repair of replication forks stalled by a nickase (34), suggesting that reversed fork restoration by AddAB might not be restricted to head-on conflicts.

Replication fork reversal is a highly conserved phenomenon. In human cells, reversed forks can be restored by both re-annealing and degradation of the nascent strands (20). The RECQ1 helicase has been shown to perform the re-annealing while the DNA2 and WRN nucleases degrade reversed replication forks and promote replication restart (35). This indicates that, mechanistically, reversed fork restoration is conserved across the tree of life.

In the past, we observed different genetic requirements for the repair of head-on conflicts, when using an engineered conflict with the *lacZ* gene. However, this construct was continuously expressed at high levels in the head-on orientation; in that context, RecA promotes survival while AddAB has a minor role (13). This is in contrast to our system, in which we can turn the conflict on by inducing transcription.

There is functional enrichment for stress response genes in the head-on orientation (36, 37). In contrast to our previous conflict system mentioned above (constitutive expression), the system used in this study, where we induced the conflict is more similar to a cellular response to stress. Many head-on genes are commonly not expressed or expressed at low levels until they are induced in response to changes in the environment (11).

The requirement of the resection machinery for replication-transcription conflict resolution has been observed in *E. coli* (6, 38, 39). One of these studies proposed that head-on conflicts lead to replication fork reversal (6). Another study proposed whole replicon degradation and fork reset (38). Our data supports the first model as the exonuclease activities of AddAB are not required for conflict resolution. Moreover, if AddAB runs into a Chi sequence in this context, it would lead to the formation of long stretches of ssDNA, which can be detrimental for cell survival. In our model, the annealed nascent strands are short, and therefore unlikely to have a Chi sequence, which allows for dsDNA degradation. We cannot exclude the mechanism of fork processing is different in *B. subtilis* compared to *E. coli*.

Notably, we have shown in the past that AddAB does not promote mutagenesis due to head-on replication-transcription conflicts (14), suggesting that replication fork reversal and subsequent re-annealing is an error-free mechanism that suppresses DNA repair pathways that promote transcription-dependent mutagenesis, specifically, transcription-coupled repair (40, 41).

Overall, we provide evidence that, upon encountering a highly transcribed gene in the head-on orientation, two DNA structures are formed. On the transcription side, the annealing of the mRNA to its template DNA forms an R-loop (11). Meanwhile, on the replication side, forks can reverse, creating create four-way junctions. It is the resolution of both structures, the R-loop by RNase HIII and the reversed fork by AddAB, that allow replication to continue and cells to overcome head-on replication-transcription conflicts.

## Materials and Methods

### Bacterial culture

*Bacillus subtilis* was cultured in lysogeny broth (LB). Bacterial plates were grown overnight at 37 °C with the following antibiotics when appropriate: 100 µg/ml spectinomycin, 500 µg/ml erythromycin and 12.5 µg/ml lincomycin (MLS), 5 µg/ml (*B. subtilis*) or 50 µg/ml (*E. coli*) kanamycin, 10 µg/ml chloramphenicol and 100 µg/ml carbenicillin. Liquid cultures were started from single colonies and grown with aeration (260 rpm). A list of all strains used in this study can be found in Table S1 and a list of plasmids can be found in Table S2.

### Strain and plasmid construction

The parental strain for all *B. subtilis* strains used in this study is HM1 (same as AG174, originally named JH642) (42, 43). Gene deletions that are marked with MLS or kanamycin resistance were obtained from (44), and transformed into HM1 as in (45) except for Δ*addAB*, which was made in the HM1 background (14). When indicated, antibiotic resistant cassettes were excised by transforming the strains with a plasmid expressing the Cre recombinase (pDR244, BGSCID: ECE274) purified from Rec+ *E. coli* cells (AG1111) (46) with the QIAprep Spin Miniprep Kit (QIAGEN), generating markerless strains.

For the transposon screen, the spectinomycin resistance gene in pWX642 (22) was switched with the kanamycin resistance gene from *mfd*::kan *B. subtilis* cells (BGSCID: BKK00550) to generate pHM785. To do this, the kanamycin resistance gene, including the promoter and ribosome biding site (RBS), and pWX642 without the spectinomycin resistance gene were PCR amplified (Q5 High-Fidelity DNA Polymerase, NEB) with primers containing 20 nucleotides of overlap. The PCR products were cloned using NEBuilder® HiFi DNA Assembly Master Mix (NEB) for 1 hour at 50 °C. All primers used in this study can be found in Table S3.

For complementation experiments, the coding region of *addB* or *addA* was amplified from HM1 gDNA (Table S3) and cloned using NEBuilder® HiFi DNA Assembly Master Mix into pCAL838 (47) (Table S2). For *addB*, the PCR product included the native RBS, while the following transcription terminator was included in the reverse primer GTCCCATTCATAAGGA; for *addA*, the native transcription terminator was included in the PCR product, and the following RBS and linker were included in the forward primer: AAGGAGGTATACAT. To make the catalytic mutants, these plasmids were amplified with primers containing the appropriate mutation and 20 nt of overlap (Table S3), and circularized using NEBuilder® HiFi DNA Assembly Master Mix (Table S2). Plasmids were linearized using KpnI and transformed into competent cells containing a deletion of the appropriate gene.

For tagging *rnhC* with the SsrA degron, approximately 500 bp at the 3’ end of the gene were amplified (Table S3) and cloned EcoRI-XbaI into pGCS (26) (Table S2). These plasmids, purified from AG1111 cells, were transformed into competent *B. subtilis* cells.

For AddAB purification, the coding sequence of *addB* was amplified (Table S3) and cloned into pET28a (Thermo) (Table S3) by HiFi (NEB) to generate a 6XHis N-terminal tagged protein. The coding sequence of *addA* was amplified (Table S3) and cloned into pET21a (Thermo) (Table S3) by HiFi (NEB) to generate a 6XHis C-terminal tagged protein.

### Transposon screen

HM2195 (11) cells were transformed with pHM785 purified from AG1111 cells, plated on MLS, and incubated at 30 °C for 40 hours. For each biological replicate (three biological replicates were done in parallel), four colonies were inoculated into four tubes with 2 ml of LB supplemented with 5 µg/ml of kanamycin, and were grown for 22 hours at 25 °C with aeration. The four cultures were mixed thoroughly and 100 µl of cells were mixed with 100 µl of 50% glycerol and frozen at -80 °C. The next day, the frozen cells were thawed in a 37 °C water bath, added to 50 ml of LB supplemented with 5 µg/ml of kanamycin, and grown for 5 hours at 37 °C with aeration. 500 µl of these cells were added to two 50 ml flasks with LB supplemented with 5 µg/ml of kanamycin, one of which had 1 mM IPTG (conflict ON condition) while the other did not (conflict OFF condition). These cultures were grown to an OD600 of approximately 0.5, after which they were spun down, and the pellets were frozen at -20 °C. The pellets were then resuspended in 1 ml of solution A (10 mM Tris–HCl pH 8.0, 20% w/v sucrose, 50 mM NaCl, 10 mM EDTA, 1 mg/ml lysozyme) and incubated at 37 °C for 30 mins. Then, cells were lysed with 1 ml lysis buffer (100 mM Tris pH 7.0, 10 mM EDTA, 2% triton X-100, 300 mM NaCl) supplemented with 40 µg/ml of RNase A, DNase and protease-free (Thermo) and incubated for 30 mins at room temperature, followed by addition of SDS to a concentration of 0.25% and 0.4 mg/ml of proteinase K, recombinant, PCR grade (Thermo) and incubation at 37 °C for 30 mins. The genomic DNA was then purified by standard phenol:chloroform extraction and ethanol precipitation, and resuspended overnight with 300 µl of TE buffer pH=8. 10 ug of gDNA were digested with 6 units of MmeI (NEB) for 3 hours at 37 °C, after which 10 units of Quick CIP (NEB) were added, and the DNA was incubated for another hour at 37 °C. The digested DNA was purified using the GeneJET PCR Purification Kit (Thermo) and ligated to a DNA duplex made of HM7071 and HM7039 (Table S3) for 16 hours at 16 °C using 400 units of T4 DNA ligase (NEB). The ligation products were cleaned with AMPure XP Beads (Beckman Coulter), and 900 ng of DNA were PCR amplified using HM7115 and HM7079 as primers for 15 cycles (Table S3). The PCR products were cleaned with AMPure XP Beads and one fourth of the eluted product was PCR amplified using an i7 and an i5 primer from the NEBNext Multiplex Oligos for Illumina (Dual Index Primers Set 2) (NEB) for 6 cycles. The PCR products were pooled, purified using the GeneJET PCR Purification Kit (Thermo) and submitted for sequencing in a NovaSeq XP Plus (2x150 flow cell) at the Vanderbilt Technologies for Advanced Genomics (VANTAGE), obtaining an average of 9.7 million reads per library.

For sequencing analysis, low quality treads and bases were trimmed with fastp (48), and the resulting reads were aligned to the HM1 genome using bowtie2 (49). Using a custom R script, all unique alignments that either start or end with TA (the targe sequence of the himar1 transposase) were obtained and the coordinate of the start of the read was assigned to the corresponding gene or non-coding region.

### Survival assays

A single colony of the indicated strain was inoculated into 2 ml of LB and grown at 37 °C with aeration to an OD600 of >0.5. This culture was then diluted to an OD600 of 0.3, and then 10-fold serially diluted in 1x Spizizen’s Salts (15 mM ammonium sulfate, 80 mM dibasic potassium phosphate, 44 mM monobasic potassium phosphate, 3.4 mM trisodium citrate, and 0.8 mM magnesium sulfate). 5 ul of each dilution was plated onto LB plates with the indicated concentration of IPTG and incubated at 30°C overnight. Plates were imaged with a BioRad Gel Doc XR+ Molecular Imager and colonies were counted after either 24 or 48 h of incubation.

### Genome copy number analysis

A single colony of the indicated strain was inoculated into 5 ml of LB and grown to an OD600 of >0.5. This culture was then diluted to an OD600 of 0.05 in LB supplemented with the indicated amount of IPTG and grown to an OD of approximately 0.5. The cultures were then centrifuged, washed with 1X PBS and frozen at -20 °C. The pellets were then resuspended in 1 ml of solution A at 37 °C for 30 mins. Then, cells were lysed with 1 ml lysis buffer, incubated on ice for 30 mins and sonicated as described for the ChIPs. 0.4 mg/ml of proteinase K was then added and the samples were incubated at 37 C for 30 mins. The genomic DNA was then purified by standard phenol:chloroform extraction and ethanol precipitation, and resuspended overnight with 100 µl of TE buffer pH=8. 1 µg of DNA was used to make sequencing libraries using the NEBNext Ultra II DNA Library Prep Kit for Illumina (NEB). The libraries were submitted for sequencing in a NovaSeq XP Plus (2x150 flow cell) at the Vanderbilt Technologies for Advanced Genomics (VANTAGE), obtaining an average of 4.5 million reads per library. Low quality treads and bases were trimmed with fastp (48), and the resulting reads were aligned to the appropriate strain using bowtie2 (49).

### AddAB purification

Both pHM803 and pHM804 were transformed together into BL21(DE3)pLysS Competent Cells (Agilent), and a single colony was inoculated into 5 ml of LB and grown overnight in LB containing kanamycin and carbenicillin. 2 ml of culture were then inoculated in 1 L of LB containing kanamycin and carbenicillin and grown until an OD600 of 0.3, when 1 mM IPTG was added to the media and the cells were moved to 25 °C, were the cells were grown for an additional 3 hours, centrifuged for 15 mins at 6000G, and resuspended in 10 ml of 50 mM Tris (pH 7.5), 10% sucrose and stored at −80 °C. 10 ml of CelLytic B cell lysis reagent (Sigma) with 2 µl of Benzonase (Sigma), 10 mM imidazole and one cOmplete™, EDTA-free Protease Inhibitor Cocktail tablet (Millipore) were added and the mixture shaken gently at RT for 10 mins.

The lysate was centrifuged at 20000G at 4 °C for 30 minutes, the supernatant was mixed with an equal volume of equilibration buffer (20 mM sodium phosphate pH 7.4, 300 mM sodium chloride, 10 mM imidazole), and run twice through 10 ml of equilibrated HisPur™ Ni-NTA Resin (Thermo) at 4 °C. The resin was washed with 50 ml of wash buffer (20 mM sodium phosphate pH 7.4, 300 mM sodium chloride, 40 mM imidazole) and eluted with 10 ml of elution buffer (20 mM sodium phosphate pH 7.4, 300 mM sodium chloride, 250 mM imidazole). The protein was dialyzed with a 15 ml Slide-A-Lyzer G3 Dialysis Cassette G2 20000 MWCO (Thermo) against 10 mM tris pH 8, 50 mM NaCl, 5% glycerol, 0.1 mM DTT, 0.1 mM EDTA overnight at 4 °C and concentrated with two Amicon Ultra-4 Centrifugal Filter Units 100000 NMWL (Millipore) to a final concentration of 0.9 mg/ml measured by Qubit (Thermo).

### AddAB digestion assay

AddAB digestion was tested in 10 µl reactions containing 1X rCutSmart buffer (NEB), 1 mM ATP (when indicated), 1.5 nM DNA substrate and 1.5 µM ET SSB (NEB). The reactions were started by adding the indicated concentration of AddAB and they were incubated at 37 °C for 10 minutes, after which they were stopped with 10 μL of 98% formamide 10 mM EDTA. DNA was denatured at 95 °C for 5 min and ran in a 10% Mini-PROTEAN® TBE-Urea Gel, 10 well, 30 µl (Biorad). Gels were scanned in a ChemiDoc imaging system (BioRad). A ladder with three Cy5 labeled 46 nt, 29 nt, and 20 nt oligos was included in all gels.

## Acknowledgements

This work was supported by K99ES037493 to JC-G and Vanderbilt University School of Medicine Department of Biochemistry startup funds to HM.

This research was also supported by the Vanderbilt School of Medicine Basic Sciences, Department of Biochemistry, Destination Biochemistry Advanced Postdoctoral Scholar Award funded by the Armstrong Family to JC-G.

## Author Contributions

J.C.-G. and H.M. designed research; J.C.- G performed research; J.C.-G. and H.M. analyzed data; and J.C.-G. and H.M. wrote the paper.

---

**SI Figure 1.**
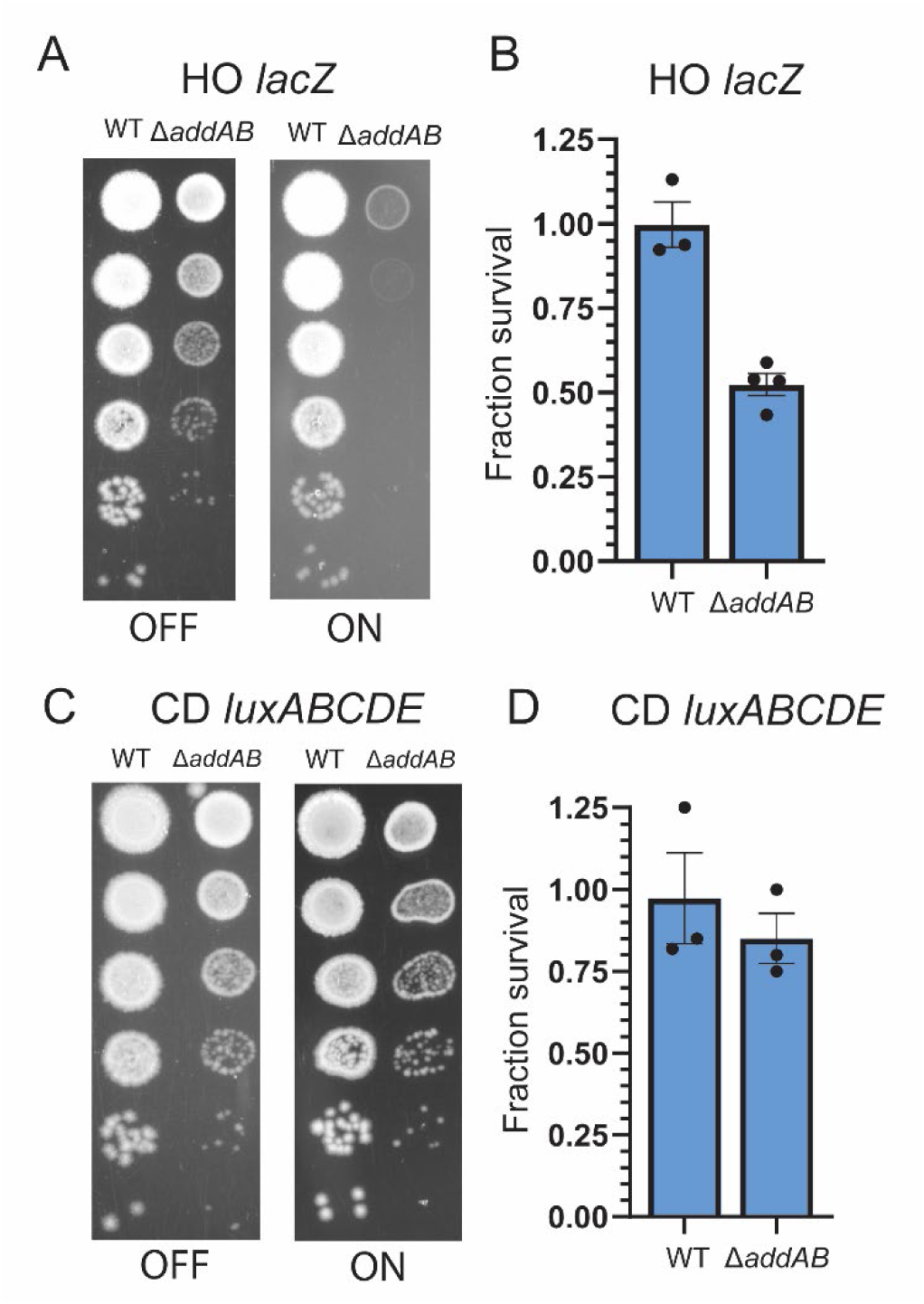
A-B) Survival assay of cells of the indicated genotype with the *lacZ* conflict integrated into *thrC* in the head-on orientation. C-D) Survival assay of cells of the indicated genotype with the *luxABCDE* conflict integrated into *amyE* in the co-directional orientation. Fraction survival was calculated as in Fig. 2C.

**SI Figure 2.**
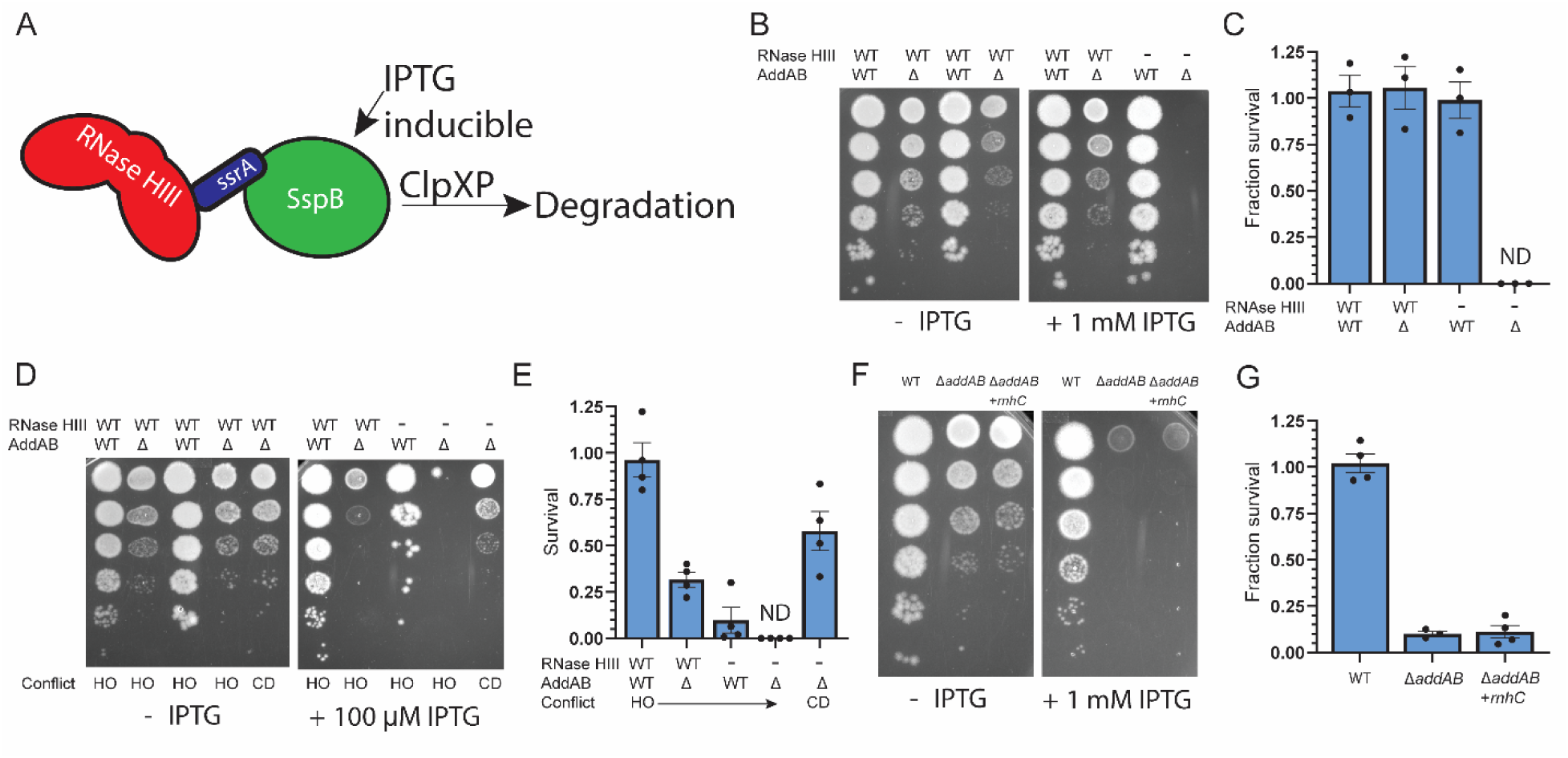
A) Schematic of the degron system used to deplete RNase III levels. A C-terminus SsrA tag placed on RNase HIII is recognized by SspB, whose expression can be induced by IPTG, and targets RNase HIII for ClpXP-dependent degradation. B-C) Survival assay of cells of the indicated genotype. D-E) Survival assay of cells of the indicated genotype. HO: Head-On conflict; CD: Co-Directional conflict. F) Survival assay of cells of the indicated genotype. +*rnhC* indicates the cells have two copies of the *rnhC* gene. Fraction survival was calculated as in Fig. 2C.

**SI Figure 3.**
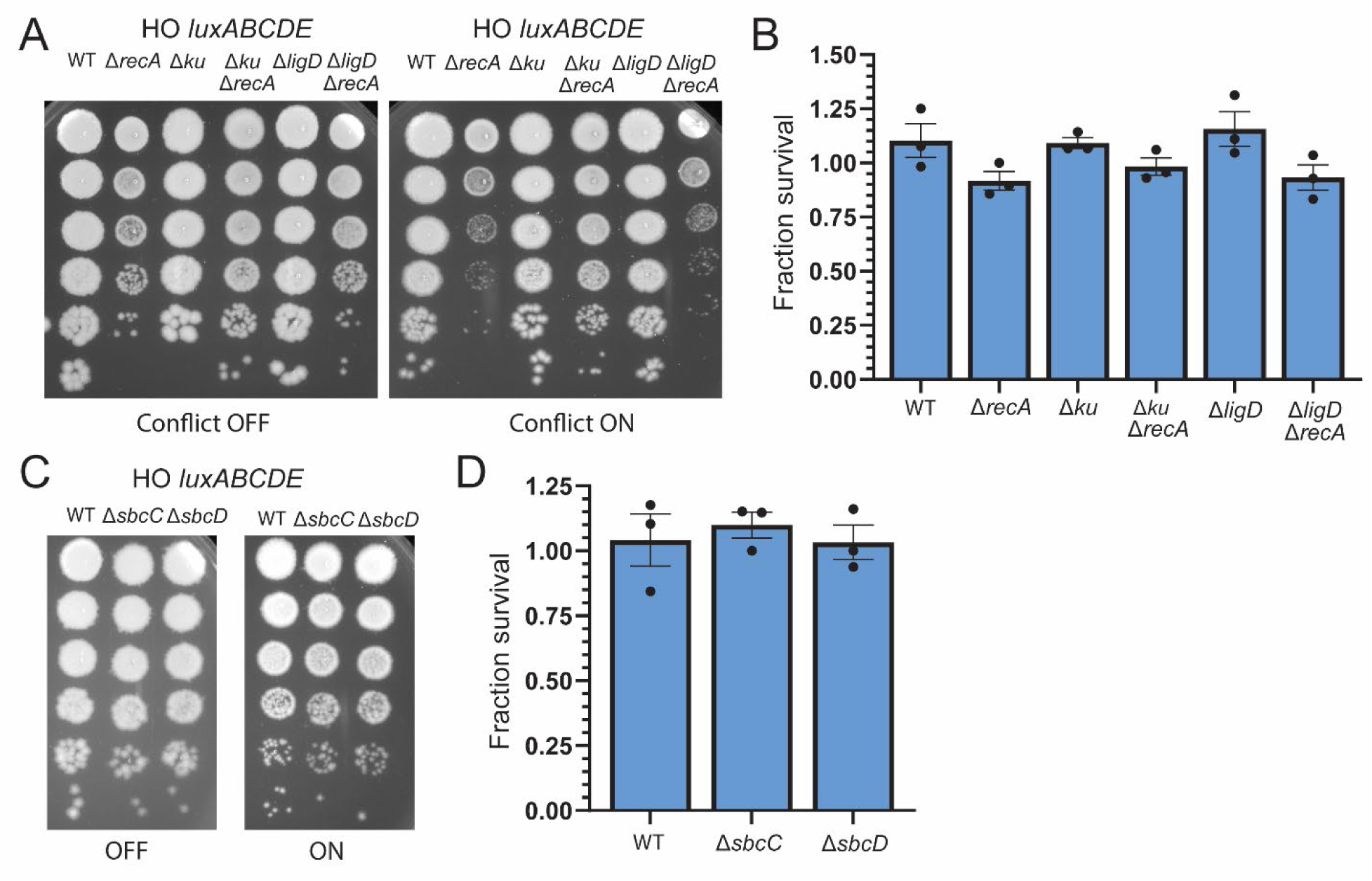
A-B) Survival assay of cells of the indicated genotype with the *luxABCDE* conflict integrated into *amyE* in the head-on orientation. C-D) Survival assay of cells of the indicated genotype with the *luxABCDE* conflict integrated into *amyE* in the head-on orientation. Fraction survival was calculated as in Fig. 2C.

**SI Figure 4.**
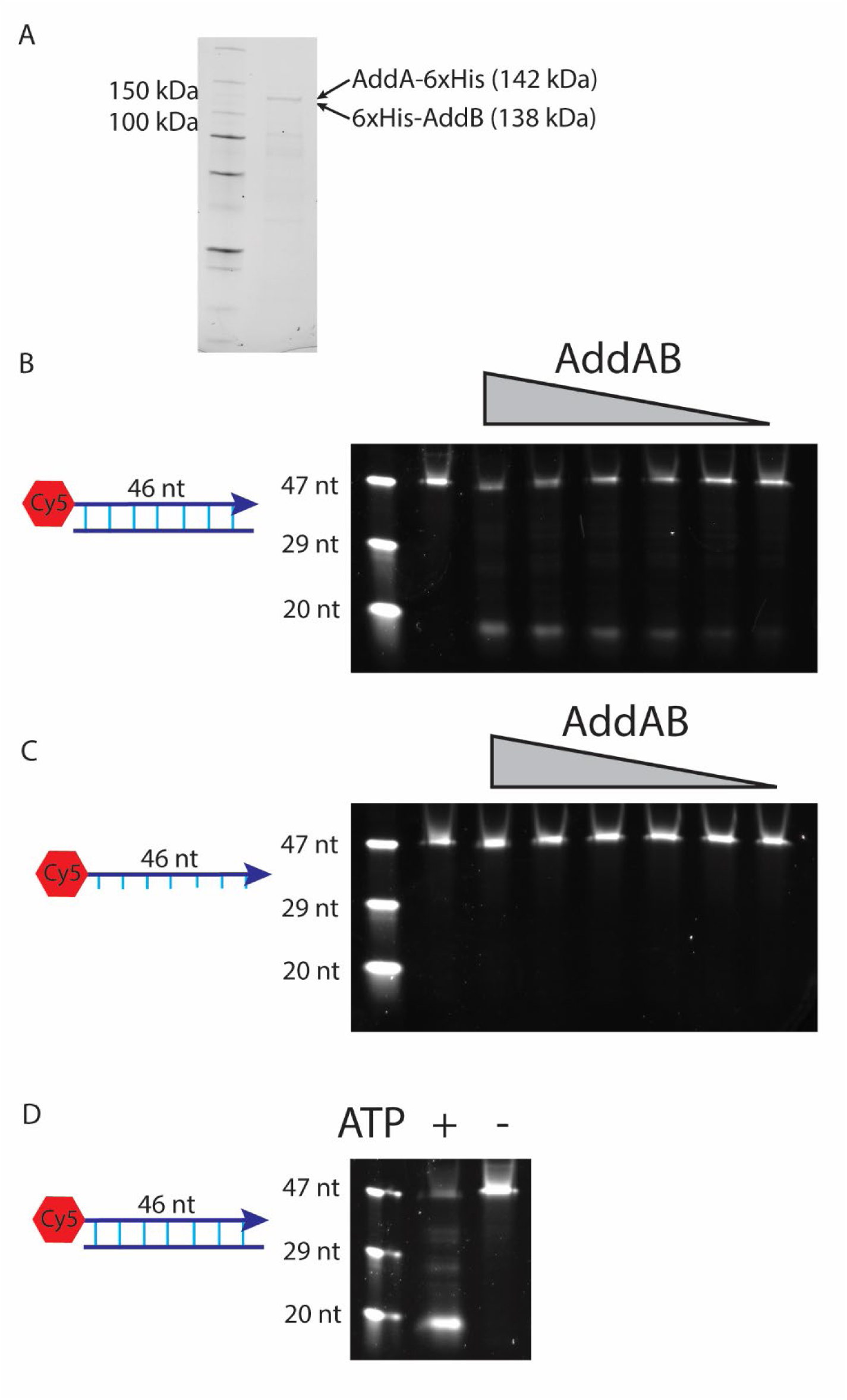
A) Purified AddAB ran in a 4-20% SDS gel imaged by Stain-Free™. B-C) 10% Urea gels of a dsDNA substrate (B) or a ssDNA substrate (C) digestion with AddAB. The highest concentration of protein is 150 nM and the lanes to the right are 2-fold dilutions. D) 10% Urea gels of the dsDNA substrate digested with 150 nM AddAB. 1 mM ATP was added when indicated.

**SI Table 1:** Strains used in this study

| Strain Name | Genotype | Figure | Reference |
| --- | --- | --- | --- |
| HM2195 | *amyE::Pspank(hy)-luxABCDE-HO* | 1,2,3,5,S3 | (11) |
| HM5032 | *amyE::Pspank(hy)-luxABCDE-HO ΔaddA::markerless* | 2 | This study |
| HM5043 | *amyE::Pspank(hy)-luxABCDE-HO ΔaddA::markerless*<br>*thrC::Pspank(hy)-addA* | 2 | This study |
| HM5013 | *amyE::Pspank(hy)-luxABCDE-HO ΔaddB::markerless* | 2 | This study |
| HM5038 | *amyE::Pspank(hy)-luxABCDE-HO ΔaddB::markerless*<br>*thrC::Pspank(hy)-addB* | 2 | This study |
| HM4985 | *amyE::Pspank(hy)-luxABCDE-HO ΔaddAB::kan* | 2,5 | This study |
| HM4991 | *amyE::Pspank(hy)-luxABCDE-HO ΔrecA::kan* | 3,S3 | This study |
| HM5117 | *amyE::Pspank(hy)-luxABCDE-HO ΔaddB::markerless*<br>*thrC::Pspank(hy)-addB-D961A* | 5 | This study |
| HM5118 | *amyE::Pspank(hy)-luxABCDE-HO ΔaddA::markerless*<br>*thrC::Pspank(hy)-addA-D1172A* | 5 | This study |
| HM5119 | *amyE::Pspank(hy)-luxABCDE-HO ΔaddA::markerless*<br>*thrC::Pspank(hy)-addA-K36A* | 5 | This study |
| HM1344 | *thrC::Pspank(hy)-lacZ-HO* | S1 | (50) |
| HM5001 | *thrC::Pspank(hy)-lacZ-HO ΔaddAB::kan* | S1 | This study |
| HM2197 | *amyE::Pspank(hy)-luxABCDE-CD* | S1 | (11) |
| HM5099 | *amyE::Pspank(hy)-luxABCDE-HO ΔaddAB::kan* | S1 | This study |
| HM447 | *thrC::Pspank-sspB* | S2 | (26) |
| HM5111 | *thrC::Pspank-sspB ΔaddAB::kan* | S2 | This study |
| HM5020 | *thrC::Pspank-sspB rnhC-ssrA* | S2 | This study |
| HM5030 | *thrC::Pspank-sspB rnhC-ssrA ΔaddAB::kan* | S2 | This study |
| HM5101 | *amyE::Pspank(hy)-luxABCDE-HO thrC::Pspank-sspB* | S2 | This study |
| HM5102 | *amyE::Pspank(hy)-luxABCDE-HO thrC::Pspank-sspB* | S2 | This study |
| HM5006 | *amyE::Pspank(hy)-luxABCDE-HO thrC::Pspank-sspB*<br>*rnhC-ssrA* | S2 | This study |
| HM5018 | *amyE::Pspank(hy)-luxABCDE-HO thrC::Pspank-sspB*<br>*rnhC-ssrA ΔaddAB::kan* | S2 | This study |
| HM4995 | *amyE::Pspank(hy)-luxABCDE-CD thrC::Pspank(hy)-*<br>*rnhC ΔaddAB::kan* | S2 | This study |
| HM5113 | *amyE::Pspank(hy)-luxABCDE-CD thrC::Pspank-sspB*<br>*rnhC-ssrA ΔaddAB::kan* | S2 | This study |
| HM5124 | *amyE::Pspank(hy)-luxABCDE-HO Δku::kan* | S3 | This study |
| HM5125 | *amyE::Pspank(hy)-luxABCDE-HO Δku::kan*<br>*ΔrecA::MLS* | S3 | This study |
| HM5126 | *amyE::Pspank(hy)-luxABCDE-HO ΔligD::kan* | S3 | This study |
| HM5127 | *amyE::Pspank(hy)-luxABCDE-HO ΔligD::kan*<br>*ΔrecA::MLS* | S3 | This study |
| HM5128 | *amyE::Pspank(hy)-luxABCDE-HO ΔsbcC::MLS* | S3 | This study |
| HM5129 | *amyE::Pspank(hy)-luxABCDE-HO ΔsbcD::MLS* | S3 | This study |

**Table S2:** Plasmids generated in this study.

| Name | Description |
| --- | --- |
| pHM785 | pWX642-kan |
| pHM796 | pCAL838-*addA* |
| pHM792 | pCAL838-*addB* |
| pHM341 | pGCS-*rnhC* |
| pHM803 | pET28a-*addB* |
| pHM804 | pET21a-*addA* |
| pHM805 | pCAL838-*addB*-D961A |
| pHM806 | pCAL838-*addA*-D1172A |
| pHM807 | pCAL838-*addA*-K36A |

**Table S3:**
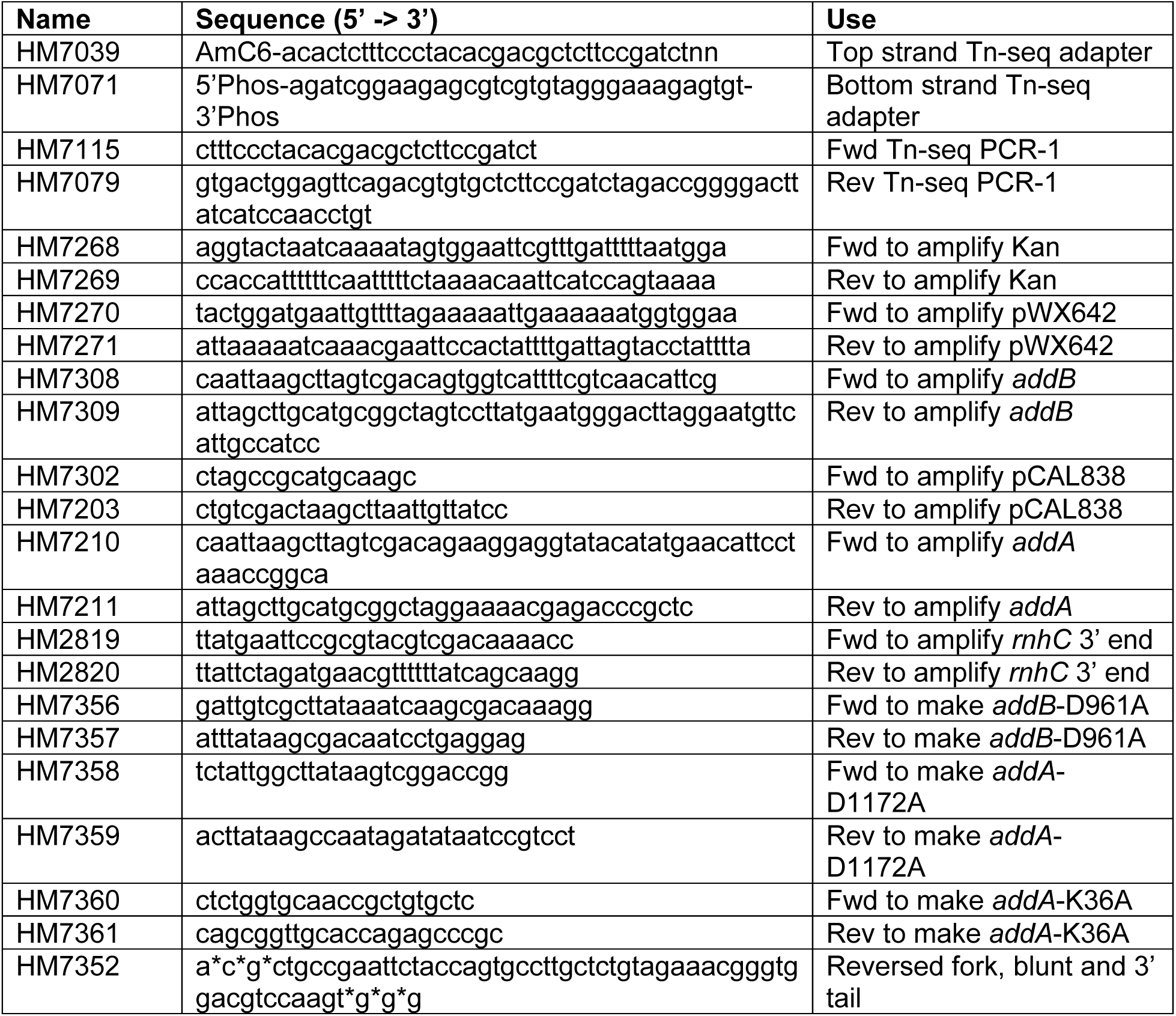
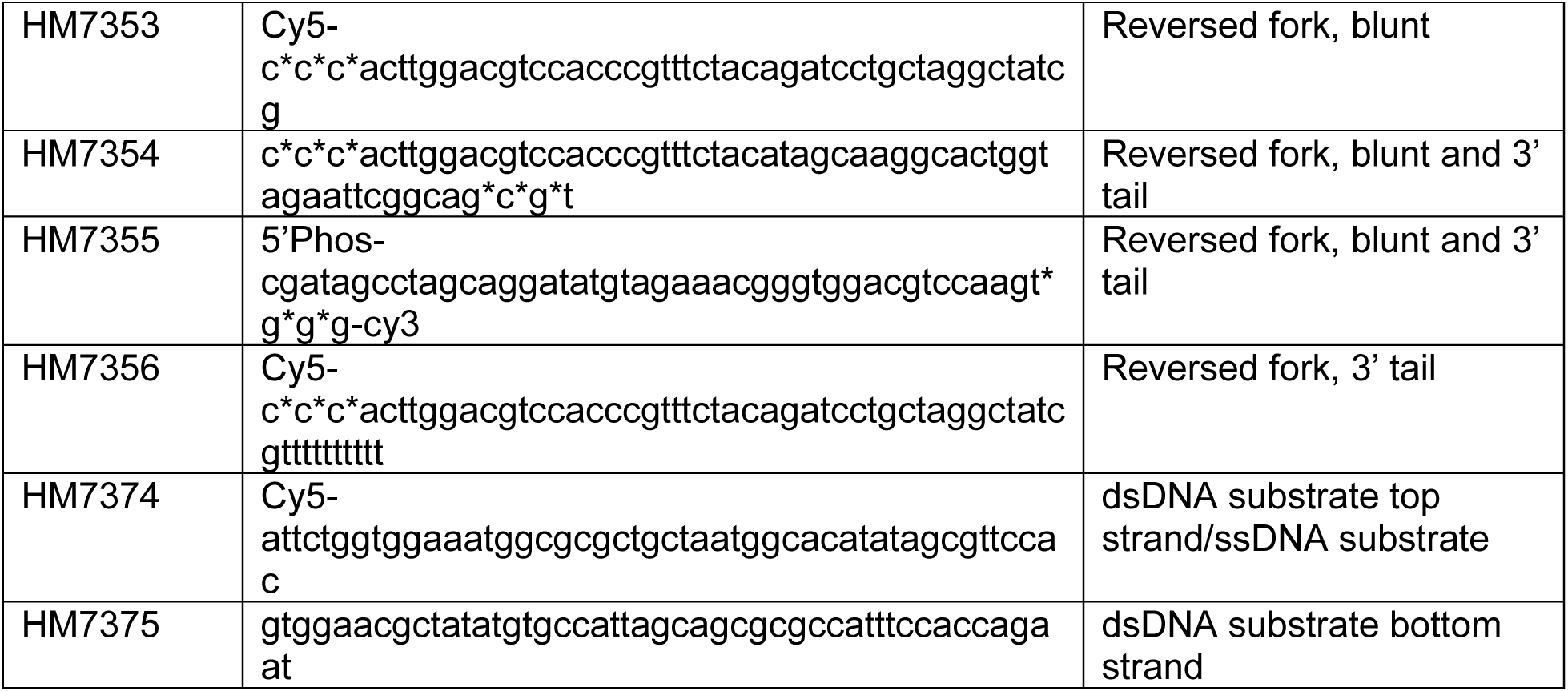
Primers used in this study

## References

1. K. R. Browning, H. Merrikh, Replication–Transcription Conflicts: A Perpetual War on the Chromosome. Annu. Rev. Biochem. 93, 21–46 (2024).

2. B. Liu, B. M. Alberts, Head-On Collision Between a DNA Replication Apparatus and RNA Polymerase Transcription Complex. Science (1979). 267, 1131–1137 (1995).

3. S. M. Mangiameli, C. N. Merrikh, P. A. Wiggins, H. Merrikh, Transcription leads to pervasive replisome instability in bacteria. Elife 6 (2017).

4. F. Prado, A. Aguilera, Impairment of replication fork progression mediates RNA polII transcription-associated recombination. EMBO J. 24, 1267–1276 (2005).

5. S. Hamperl, M. J. Bocek, J. C. Saldivar, T. Swigut, K. A. Cimprich, Transcription-Replication Conflict Orientation Modulates R-Loop Levels and Activates Distinct DNA Damage Responses. Cell 170, 774–786.e19 (2017).

6. A. L. De Septenville, S. Duigou, H. Boubakri, B. Michel, Replication Fork Reversal after Replication–Transcription Collision. PLoS Genet. 8, e1002622 (2012).

7. S. Paul, S. Million-Weaver, S. Chattopadhyay, E. Sokurenko, H. Merrikh, Accelerated gene evolution through replication–transcription conflicts. Nature 495, 512–515 (2013).

8. T. S. Sankar, B. D. Wastuwidyaningtyas, Y. Dong, S. A. Lewis, J. D. Wang, The nature of mutations induced by replication–transcription collisions. Nature 535, 178–181 (2016).

9. E. P. C. Rocha, The replication-related organization of bacterial genomes. Microbiology (N. Y). 150, 1609–1627 (2004).

10. Y.-H. Chen, et al., Transcription shapes DNA replication initiation and termination in human cells. Nat. Struct. Mol. Biol. 26, 67–77 (2019).

11. K. S. Lang, et al., Replication-Transcription Conflicts Generate R-Loops that Orchestrate Bacterial Stress Survival and Pathogenesis. Cell 170, 787–799.e18 (2017).

12. P. Huertas, A. Aguilera, Cotranscriptionally Formed DNA:RNA Hybrids Mediate Transcription Elongation Impairment and Transcription-Associated Recombination. Mol. Cell 12, 711–721 (2003).

13. S. Million-Weaver, A. N. Samadpour, H. Merrikh, Replication Restart after Replication-Transcription Conflicts Requires RecA in Bacillus subtilis. J. Bacteriol. 197, 2374–2382 (2015).

14. S. Million-Weaver, et al., An underlying mechanism for the increased mutagenesis of lagging-strand genes in Bacillus subtilis. Proc. Natl. Acad. Sci. U. S. A. 112, E1096–E1105 (2015).

15. J. Sollier, et al., Transcription-Coupled Nucleotide Excision Repair Factors Promote R-Loop-Induced Genome Instability. Mol. Cell 56, 777–785 (2014).

16. K. S. Lang, H. Merrikh, Topological stress is responsible for the detrimental outcomes of head-on replication-transcription conflicts. Cell Rep. 34, 108797 (2021).

17. R. E. Wellinger, F. Prado, A. Aguilera, Replication Fork Progression Is Impaired by Transcription in Hyperrecombinant Yeast Cells Lacking a Functional THO Complex. Mol. Cell. Biol. 26, 3327–3334 (2006).

18. D. B. Wigley, Bacterial DNA repair: recent insights into the mechanism of RecBCD, AddAB and AdnAB. Nat. Rev. Microbiol. 11, 9–13 (2013).

19. F. Chédin, S. D. Ehrlich, S. C. Kowalczykowski, The Bacillus subtilis AddAB helicase/nuclease is regulated by its cognate Chi sequence in vitro. J. Mol. Biol. 298, 7–20 (2000).

20. K. J. Neelsen, M. Lopes, Replication fork reversal in eukaryotes: from dead end to dynamic response. Nat. Rev. Mol. Cell Biol. 16, 207–220 (2015).

21. C. N. Merrikh, B. J. Brewer, H. Merrikh, The B. subtilis Accessory Helicase PcrA Facilitates DNA Replication through Transcription Units. PLoS Genet. 11, e1005289 (2015).

22. G. S. Dobihal, J. Flores-Kim, I. J. Roney, X. Wang, D. Z. Rudner, The WalR-WalK Signaling Pathway Modulates the Activities of both CwlO and LytE through Control of the Peptidoglycan Deacetylase PdaC in Bacillus subtilis. J. Bacteriol. 204 (2022).

23. S. Sanchez, E. V Snider, X. Wang, D. B. Kearns, Identification of Genes Required for Swarming Motility in Bacillus subtilis Using Transposon Mutagenesis and High-Throughput Sequencing (TnSeq). J. Bacteriol. 204, e0008922 (2022).

24. J. T. P. Yeeles, E. J. Gwynn, M. R. Webb, M. S. Dillingham, The AddAB helicase–nuclease catalyses rapid and processive DNA unwinding using a single Superfamily 1A motor domain. Nucleic Acids Res. 39, 2271–2285 (2011).

25. M. Moreno-del Álamo, B. Carrasco, R. Torres, J. C. Alonso, Bacillus subtilis PcrA Helicase Removes Trafficking Barriers. Cells 10, 935 (2021).

26. K. L. Griffith, A. D. Grossman, Inducible protein degradation in *Bacillus subtilis* using heterologous peptide tags and adaptor proteins to target substrates to the protease ClpXP. Mol. Microbiol. 70, 1012–1025 (2008).

27. S. Shuman, M. S. Glickman, Bacterial DNA repair by non-homologous end joining. Nat. Rev. Microbiol. 5, 852–861 (2007).

28. B. M. Wendel, J. M. Cole, C. T. Courcelle, J. Courcelle, SbcC-SbcD and ExoI process convergent forks to complete chromosome replication. Proceedings of the National Academy of Sciences 115, 349–354 (2018).

29. H. Stoy, et al., Direct visualization of transcription-replication conflicts reveals post-replicative DNA:RNA hybrids. Nat. Struct. Mol. Biol. 30, 348–359 (2023).

30. J. T. P. Yeeles, M. S. Dillingham, A Dual-nuclease Mechanism for DNA Break Processing by AddAB-type Helicase-nucleases. J. Mol. Biol. 371, 66–78 (2007).

31. F. Chédin, P. Noirot, V. Biaudet, S. D. Ehrlich, A five-nucleotide sequence protects DNA from exonucleolytic degradation by AddAB, the RecBCD analogue of *Bacillus subtilis*. Mol. Microbiol. 29, 1369–1377 (1998).

32. M. Berti, et al., Human RECQ1 promotes restart of replication forks reversed by DNA topoisomerase I inhibition. Nat. Struct. Mol. Biol. 20, 347–354 (2013).

33. N. Kim, S. Jinks-Robertson, Transcription as a source of genome instability. Nat. Rev. Genet. 13, 204–214 (2012).

34. C. Winterhalter, K. J. Stratton, S. Fenyk, H. Murray, Rescuing the bacterial replisome at a nick requires recombinational repair and helicase reloading. Nat. Commun. 16, 11633 (2025).

35. S. Thangavel, et al., DNA2 drives processing and restart of reversed replication forks in human cells. Journal of Cell Biology 208, 545–562 (2015).

36. S. Paul, S. Million-Weaver, S. Chattopadhyay, E. Sokurenko, H. Merrikh, Accelerated gene evolution through replication–transcription conflicts. Nature 495, 512–515 (2013).

37. C. N. Merrikh, H. Merrikh, Gene inversion potentiates bacterial evolvability and virulence. Nat. Commun. 9, 4662 (2018).

38. M. B. Cooke, et al., Transcription–replication collisions trigger high-fidelity replication reset. Nucleic Acids Res. 53 (2025).

39. E. A. Kouzminova, G. E. Cronan, A. Kuzminov, RecBCD-dependent post-UV replication restart in Escherichia coli triggers fork triplication. Proceedings of the National Academy of Sciences 123 (2026).

40. M. N. Ragheb, et al., Inhibiting the Evolution of Antibiotic Resistance. Mol. Cell 73, 157–165.e5 (2019).

41. J. Carvajal-Garcia, A. N. Samadpour, A. J. Hernandez Viera, H. Merrikh, Oxidative stress drives mutagenesis through transcription-coupled repair in bacteria. Proceedings of the National Academy of Sciences 120 (2023).

42. S. P. Brehm, S. P. Staal, J. A. Hoch, Phenotypes of Pleiotropic-Negative Sporulation Mutants of *Bacillus subtilis*. J. Bacteriol. 115, 1063–1070 (1973).

43. J. L. Smith, J. M. Goldberg, A. D. Grossman, Complete Genome Sequences of Bacillus subtilis subsp. *subtilis* Laboratory Strains JH642 (AG174) and AG1839. Genome Announc. 2 (2014).

44. B.-M. Koo, et al., Construction and Analysis of Two Genome-Scale Deletion Libraries for Bacillus subtilis. Cell Syst. 4, 291–305.e7 (2017).

45. S. M. C. C. R. Harwood, Molecular biological methods for Bacillus. Wiley (1990).

46. K. Ireton, A. D. Grossman, Coupling between gene expression and DNA synthesis early during development in Bacillus subtilis. Proceedings of the National Academy of Sciences 89, 8808–8812 (1992).

47. C. T. Leonetti, et al., Critical Components of the Conjugation Machinery of the Integrative and Conjugative Element ICE *Bs1* of Bacillus subtilis. J. Bacteriol. 197, 2558–2567 (2015).

48. S. Chen, fastp 1.0: An ultra-fast all-round tool for FASTQ data quality control and preprocessing. iMeta 4 (2025).

49. B. Langmead, S. L. Salzberg, Fast gapped-read alignment with Bowtie 2. Nat. Methods 9, 357–359 (2012).

50. O. Sensoy, J. Carvajal-Garcia, J. R. Heyza, P. A. Wiggins, H. Merrikh, A universal regulatory mechanism for prevention of replication restart from RNA:DNA hybrids. [Preprint] (2026).

